# *Fusarium oxysporum* f. sp. vasinfectum race 4 (Fov4) *FNP1,* a unique non-ribosomal peptide synthetase gene, plays an important role in fungal development and cotton Fusarium wilt pathogenesis

**DOI:** 10.1101/2025.08.11.669693

**Authors:** Huan Zhang, Yi Zhou, Caleb Oliver Bedsole, Won Bo Shim

**Affiliations:** Department of Plant Pathology and Microbiology, Texas A&M University, College Station, TX 77843, USA

**Keywords:** *Fusarium oxysporum* f. sp. *vasinfectum*, NRPS, pathogenicity, virulence, fusaric acid

## Abstract

Fusarium wilt, caused by *Fusarium oxysporum* f. sp. *vasinfectum* (Fov), is one of the most destructive early-season cotton diseases worldwide. The recent emergence of the highly virulent Fov race 4 (Fov4) and its aggressiveness have raised significant concerns for the U.S. cotton industry. Unlike predominant Fov races in US cotton production, which require root-knot nematodes to cause damage, Fov4 is known to infect cotton independent of nematodes.

However, molecular mechanisms of Fov4 virulence in cotton are not clearly understood. Secondary metabolites are often identified as the culprits in pathogen virulence toward plant hosts. To investigate these factors in Fov4, we analyzed the genomes of Fov1 and Fov4 using Fungal antiSMASH and identified a Fov4-specific nonribosomal peptide synthetase (NRPS) gene *FNP1*. To investigate its function, we generated *FNP1* knock-out mutant using CRISPR-Cas9 approach. Growth assays revealed that the mutants exhibit significantly attenuated hyphal production on media containing cotton roots as the sole carbon source, increased sensitivity to cell stress agents, as well as lagged spore germination. Furthermore, the mutant exhibited defect in cotton root rot virulence and significant decrease in Fusaric acid production. Microscopic observation of GFP-labeled *FNP1* deletion mutant showed impeded infection progression in cotton roots compared to the wild type (WT), which further explained the impeded virulence in *FNP1* mutant. Gene complementation restored the observed defects, confirming that *FNP1* is critical for Fov4 virulence, hyphal development, Fusaric acid production, and stress responses.

**Highlights:**

Comparative genomic analysis between Fov1 and Fov4 identified *FNP1* as a gene specific to Fov4.
CRISPR/Cas9 system was employed in Fov4 to generate gene deletion mutants and GFP labeling.
*FNP1* plays a critical role in Fov4 hyphal development, virulence, fusaric acid production, and stress responses.
This study is the first report to identify and functionally characterize a virulence gene in *Fusarium oxysporum* f. sp. *vasinfectum* (Fov) race 4 (Fov4) in cotton wilt pathogenesis.

## 1. Introduction

Cotton (*Gossypium spp.*) is one of the most important natural fiber crops globally, serving as a cornerstone of the textile industry and a significant contributor to agricultural economies (Khan, et al. 2020). It is primarily cultivated from four species: *Gossypium hirsutum*, *G. barbadense*, *G. arboreum*, and *G. herbaceum*. In addition to providing raw material for textiles, cotton offers valuable by-products such as cottonseed oil and meal, which are utilized in food and feed industries. Among these, *G. hirsutum* (Upland cotton) and *G. barbadense* (Pima cotton) dominate global production due to their superior fiber qualities and adaptability to diverse growing conditions (Chaudhry and Guitchounts 2003; Tausif, et al. 2018; Shahrajabian, et al. 2020). Upland cotton generally is used for medium-quality and mass-production textile products. Pima cotton is an extra-long staple cotton used in higher value textile products because of its fine and long fibers (Go, et al.). In the United States, cotton is cultivated primarily in southern states, with Texas being the leading producer, accounting for more than 40% of total U.S. production. The majority of cotton grown in the U.S. is Upland cotton which accounts for 97% of the cotton production, while Pima cotton represents about 3% of total production and is mainly grown in California, Arizona, and west Texas (USDA-ERS 2022). Despite its economic importance, cotton production faces significant challenges, particularly from biotic stresses, such as pests and diseases, that impact yield and fiber quality (Constable and Bange 2015; Hussain, et al. 2023).

One of the key early season, and perhaps most destructive, diseases of cotton is Fusarium wilt caused by *Fusarium oxysporum* f. sp *vasinfectum* (Fov) (Cianchetta, et al. 2015).

Fov is a soilborne fungal pathogen responsible for Fusarium wilt, a devastating disease that significantly impacts cotton production worldwide (Ayubov, et al. 2024). This pathogen belongs to the *F. oxysporum* species complex, which encompasses diverse strains exhibiting host-specific pathogenicity (Zhang and Ma 2017). Fov infection leads to cotton vascular wilts, damping-off, and root rot, causing severe yield losses and reducing fiber quality (Cianchetta and Davis 2015). The molecular interactions between the pathogen and host plants remain poorly understood due to challenges in applying advanced molecular genetic techniques. Investigating these interactions is essential for unraveling the mechanisms of fungal pathogenicity and developing effective management strategies. Such studies could also help address the increasing threat posed by the pathogen’s adaptability and its ability to overcome host resistance.

Historically, eight nominal Fov races are recognized as the causal agent of Fusarium wilt based on their virulence to cotton and diverse non-cotton species (Armstrong and Armstrong 1960). However, the race designations were recently revised (Davis, et al. 2006; Halpern, et al. 2020), and race 4 (Fov4) has emerged as a particularly significant threat to cotton production in the United States. First identified in California in 2001, Fov4 has since spread to major cotton-growing regions, including El Paso and Hudspeth counties in Texas and New Mexico, where it is now considered endemic (Kim, et al. 2005; Halpern, et al. 2018; Zhu, et al. 2020). Unlike earlier races, such as Fov1, which require root-knot nematodes to cause visible damage, Fov4 infects cotton independent of nematodes (Colyer, et al. 1997; El-Zik and Thaxton 1998). Its ability to thrive across diverse soil textures and pH levels, combined with the absence of resistant cotton cultivars, underscores its potential for widespread impact. Fov4 is highly virulent and affects both Pima and Upland varieties of cotton. Also its persistence in soil and host adaptability underscore the urgent need to better understand its pathogenesis mechanisms and develop integrated disease management strategies to mitigate its effects on cotton production (Ulloa, et al. 2016; Zhu 2022; Zhu, et al. 2022; Ulloa, et al. 2023).

Secondary metabolites play crucial roles in determining the virulence, pathogenicity, and host adaptability in fungal pathogens. *F. oxysporum* is known to produce a variety of mycotoxins and biologically active metabolites, including fusaric acid, heptaketides, beauvericin, and bikaverin, which can have significant effects on cotton crops (Desjardins and Proctor 2007; Ismaiel and Papenbrock 2015; Ekwomadu, et al. 2021). Secondary metabolites are primarily synthesized through non-ribosomal peptide synthetases (NRPS) and polyketide synthases (PKS), enzymes that facilitate the incorporation and enzymatic modification of building blocks to generate structurally diverse compounds (Go, et al. 2021; Lin, et al. 2023). Fusaric acid (FA), a polyketide-derived secondary metabolite, was first identified in *F. heterosporum* in 1937 and is produced by several *Fusarium* species, including *F. oxysporum* and members of the *F. fujikuroi* species complex (Bacon, et al. 1996). FA plays a key role in plant diseases such as vascular wilt, damping-off, and root rot in vegetable crops (Gaumann 1957; Stipanovic, et al. 2011). In addition to its pathogenic effects, FA is a natural contaminant in cereal grains and animal feed, presenting potential risks to agriculture and ecosystems (Smith and Sousadias 1993; Porter, et al. 1995). Although mildly toxic to mammals, FA remains a critical research focus due to its impact on plant health and its presence in agricultural commodities (Crutcher, et al. 2015; Crutcher, et al. 2017). Comparative genomic analyses have provided valuable insights into the genes and genome features involved in *Fusarium* pathogenesis, suggesting potential strategies for improved disease management (Ma, et al. 2010; Hoh, et al. 2022; Sun, et al. 2025).

In our study, we identified a NRPS gene, designated *FNP1*, that is uniquely present in Fov4 by comparing the published genomes of Fov4 and Fov1(Seo, et al. 2020). We hypothesize that *FNP1* contributes to Fov4 virulence and investigated the role of *FNP1* in Fov4 physiology and virulence. We characterized its function in vegetative growth, virulence on cotton, and fusaric acid production. Our findings provide new insights into the molecular mechanisms underlying Fov4 virulence that will help develop strategies for Fusarium wilt management.

## Materials and Methods

### 2.1 Fungal strains, culture media and nucleic acid manipulation

*Fusarium oxysporum* f. sp*. vasinfectum* race 4 (Fov4) strain used in this study was isolated from diseased Pima cotton plants collected in El Paso, Texas (Thomas Isakeit, Texas A&M AgriLife Extension). The genome of this strain was sequenced and assembled by Plasmidsaurus Inc. (Louisville, KY) using the PacBio Sequel platform. Assembly was carried out with Hifiasm (v0.16.0) and subsequently polished using Medaka (v1.8.0). The finalized genome assembly (GenBank accession number PRJNA1298872) was utilized in this study. Conidia were produced by incubating the strain in YEPD (Yeast Extract 3 g/L, Peptone 10 g/L, Dextrose 20 g/L) medium for 7 days. The fungal culture was filtered through Miracloth to collect conidia, which were then pelleted by centrifugation, resuspended, and diluted to the desired concentration.

### 1.2 Vegetative phenotypes and growth assays

For fungal growth assays, a 5-μL droplet of spore suspension (10⁵ conidia/mL) was inoculated at the center of the agar plates and incubated at 22°C. The vegetative phenotypes and growth rates (colony diameter) were assayed after 4 and 7 days post inoculation on the plates of 0.5× PDA (PDA 20 g/L, 15 g/L agar), YEPD agar (Yeast Extract 3 g/L, Peptone 10 g/L, Dextrose 20 g/L, 15 g/L agar), cotton roots modified (0.5 g/L MgSO₄·7H₂O, 1 g/L NH₄NO₃, 1 g/L KH₂PO₄, 1 g/L yeast extract, and 1 g/L cotton roots, 15 g/L agar) plates.

For the growth assay on the cotton roots only plates (1 g/L ground cotton roots, 15 g/L agar), equal volumes of conidial suspension at identical concentration were inoculated at the center of the plates. The plates were incubated at 22°C for three weeks until a clear growth phenotype developed, after which they were imaged.

### 1.3 Stress assays

For stress assays, a 5-μL droplet of conidial suspension (10⁵ /mL) was inoculated at the center of Czapek-Dox agar plates (2 g/L NaNO₃, 0.5 g/L MgSO₄·7H₂O, 0.5 g/L KCl, 10 mg/L FeSO₄·7H₂O, 1 g/L K₂HPO₄, 30 g/L sucrose, and 15 g/L agar; pH 7.3) amended with stress agents SDS (0.01%), Congo Red (50 μg/mL), or KCl (0.5 M). Plates were incubated at 22°C for 7 days.

### 1.4 Spore production and germination efficiency assay

For spore production quantification, a 5-μL droplet of conidial suspension (10⁵ /mL) was inoculated at the center of V8 agar plates (200 mL/L V8 Juice, 3g/L CaCO_3_, 20g/L agar) and incubated at 22°C for 10 days. After incubation, spores were harvested and quantified. For the spore germination efficiency assay, conidia were inoculated into 0.2× potato dextrose broth (PDB) at a concentration of 10^5^ conidia/mL and incubated at room temperature with gentle agitation (100 rpm) for 5 hours. Following incubation, 50 images were captured and germ tube lengths were measured by Image J software version 1.54f (Schneider, et al. 2012).

### 1.5 Gene deletion and gene complementation

Gene deletion mutation was carried out using a modified CRISPR/Cas9 approach based on the method developed by Wang and colleagues (Wang, et al. 2018). The EnGen® Spy Cas9 NLS (NEB, M0646M) was used in our study. Hygromycin B phosphotransferase (*HPH*) gene fragment was amplified from the pBS15 plasmid (Zhang 2017). The 5′ and 3′ flanking regions of the *FNP1* gene were fused to the *HPH* fragment on the left and right sides, respectively, by double-joint PCR. The PCR amplicons, along with sgRNA and the EnGen® Spy Cas9 NLS, were transformed into wild-type (WT) fungal protoplasts. For complementation, respective WT gene, driven by the native promoter, were co-transformed with a geneticin-resistant gene (Hoogendoorn, et al.) into mutant protoplasts. Drug-resistant colonies were subsequently screened by PCR with the Phire Plant Direct PCR Kit (Thermo Scientific, Waltham, MA, USA) (Fig. S4). To confirm the insertion of a single copy of the *HPH* gene, a droplet digital PCR (ddPCR) assay was performed using genomic DNA extracted from gene knockout mutant candidates. A duplex ddPCR assay targeting the HPH gene and the β-tubulin gene as a reference was conducted as previously described (Cai, et al. 2021). The reactions were partitioned into droplets and analyzed using the Bio-Rad AutoDG QX200 system. A resulting *HPH* to reference gene ratio of approximately 1:1 indicates a single-copy integration of the *HPH* gene into the genome. The complementation strain candidate was validated using qPCR to confirm gene expression following the complementation. All primers used in this study are listed in Table S1.

### 2.6 Generation of green fluorescence protein (GFP)-tagged strains

A fusion PCR-based method was used to generate the fragment which targets the native *FNP1* locus in the WT strain for the fluorescent strain construction. Briefly, around 1-kb upstream of the stop codon and the 3^’^ flanking region of *FNP1* (RF) were amplified. GFP was amplified from pKNTG plasmid (Yang, et al. 2018). Three fragments (1-kb upstream, GFP and RF) were fused by joint-PCR method. The joint-PCR fragments together with PBSG plasmid (geneticin-resistant marker) were transformed into WT protoplasts. This will allow GFP to be inserted to replace the *FNP1* stop codon. *FNP1* will be expressed under its native promoter and a single copy of the gene existing in the genome. The Phire plant direct PCR kit was used to confirm the correct single insertion in this study.

### 2.7 Pathogenicity assay

Upland cotton (*Gossypium hirsutum*) seeds (Ortiz, et al. 2017) were surface sterilized with 10% bleach and 70% ethanol following standard laboratory practice, and subsequently the seeds were planted in sandwich bags filled with autoclaved garden soil mix (Miracle-Gro) for 1 week in Conviron growth chamber at 23°C. After 1 week, seedlings were harvested by cutting the sandwich bag and gently removing the cotton plants. The roots were washed with tap water to remove soil, followed by three rinses with sterilized water. WT, mutant and complementation Fov4 spore suspensions (10^7^ conidia /mL) were collected from fungal cultures grown in YEPD broth and passed through a filter paper (No. 2, Whatman). The cotton roots were immersed in the spore suspension for 5 hours on a shaker at 100 rpm. Autoclaved YEPD broth was used as a mock control. After immersion, the seedlings were transplanted into glass jars containing autoclaved perlite and subsequently watered. The plants were kept in a growth chamber (23℃ and 50% RH) for 4 weeks, watering as needed. Disease evaluation was performed by visual scoring based on disease severity (percentage scale from 0 to 100% based on the root discoloration) (Merga 2018). Root surface area was measured by Image J. Fungal recovery was quantified by cutting stems after surface sterilization into approximately 1 cm pieces and incubating on PDA plates amended with kanamycin (50 µg/mL) and ampicillin (50 µg/mL).

### 2.8 Fusaric acid (FA) production assays

For FA production, WT, mutant and complementation strains were inoculated into 100 ml Czapek Dox broth and incubated at room temperature on a rotary shaker for 14 days. The cultures were then filtrated with sterile 0.2 μm pore-size vacuum filtration system (polyethersulfone, PES; Nalgene) to remove mycelia and conidia. The supernatant was adjusted to pH 2.5 with 2 M HCl and extracted with ethyl acetate three times. The combined ethyl acetate extractions were dried on a rotary evaporator, and the resultant residues were dissolved in 1 mL methanol. The samples were then analyzed by the Laboratory for Biological Mass Spectrometry (LBMS) at Texas A&M University. Three replicates were performed for each strain. Target liquid chromatography tandem mass spectrometry (LC-QQQ) analysis was performed on a TSQ Quantiva mass spectrometer (Thermo Scientific) coupled to a binary pump UHPLC (Ultimate3000, Thermo Scientific). Chromatographic separation was achieved on a Hypersil Gold 5 µm, 50 mm x 3 mm C18 column (Thermo Scientific) maintained at 30 °C using a solvent gradient method. Sample acquisition and data analysis was performed Trace Finder 3.3 (Thermo Scientific).

### 2.9 Cell imaging and staining

GFP-tagged conidia from WT and mutant were collected and used to prepare a spore suspension (10⁷ conidia/mL) in YEPD medium. Cotton roots (10-day-old) were collected and rinsed with water, dipped into the spore suspension and subsequently transferred to petri dishes containing wet filter paper. Roots were collected at 3-days post-inoculation (dpi), cut into small sections (∼2 cm) using a scalpel or razor blade, and mounted in 6% agarose either vertically or horizontally to obtain cross or longitudinal sections. These sectioned root samples were prepared using a vibratome (Leica VT1000 S, Leica Biosystems, Germany) to a thickness of 125 µm, were carefully separated from the agarose under a dissecting microscope, and stained with propidium iodide (50 µg/mL) for 10 ∼ 30 minutes. Microscopic imaging was performed using an Olympus FV3000 FluoView™ confocal laser scanning system integrated with an Olympus IX83 inverted microscope equipped with a Galvanometer scanner and high-sensitivity GaAsP PMT detectors. Two objectives were used for imaging: the UPLSAPO 20× (NA 0.75) and UPLSAPO 60× (NA 1.30) with silicone oil immersion. GFP fluorescence was excited at 488 nm and collected between 500–540 nm. Propidium iodide (PI) fluorescence was excited at 561 nm and collected between 570–670 nm. Fluorescence and transmitted light images were acquired simultaneously using FluoView software. Images were adjusted for brightness and contrast, cropped, and assembled into final figures using Adobe Photoshop version 25.12.3 (Inc. 2025).

## 3 Results

### 3.1 Gene identification

Earlier published studies suggested that Fov4 can cause Fusarium wilt without root-knot nematodes (RKN) (Kim, et al. 2005; Ayubov, et al. 2024). Also, based on the Fov genomes published by Seo et al (Seo, et al. 2020), Fov4 is predicted to harbor 2,247 more genes compared to Fov1. These earlier studies lead us to hypothesize that Fov4 employs distinct mechanisms for host recognition, infection, and penetration compared to Fov1. To investigate putative virulence factors contributing to the differences, we searched for unique genes involved in secondary metabolite biosynthesis gene clusters (BGCs) in Fov4 relative to Fov1 using antiSMASH (Blin, et al. 2021). The GenBank accession numbers VINP00000000.1 (Fov4 isolate 89-1A) and VINL00000000.1 (Fov1 isolate TF1) were used as input with default settings to initiate the analysis. Our analysis identified 26 NRPS clusters in Fov1 and 31 NRPS clusters in Fov4.

Subsequent BLASTp search comparing Fov4 NRPS sequences against Fov1 NRPS clusters identified one NRPS protein located on scaffold 12 (listed as 12.2), designated Fnp1(the first identified and studied Fov4 NRPS protein), which is specific to Fov4. It shared only 41.95% amino acid sequence identity with its closest match in Fov1, located on scaffold 10 (listed as 10.1) (Fig. S1). *FNP1* is an 8,688 bp gene predicted to encode a 2,252 amino-acid protein, featuring three conserved condensation (C) domains, one adenylation (A) domain, and two phosphopantetheine (Krappmann) binding domains, as identified by antiSMASH analysis (Fig. S2). We also identified the *FNP1* gene in our Fov4 isolate from El Paso, Texas, through BLAST analysis. This gene is 100% conserved in our isolate, as well as in eight Fov4 isolates sequenced in the Antony Babu lab at Texas A&M University (unpublished data). We concluded that *FNP1* is a specific gene in Fov4 and is absent in Fov1.

### 3.2 Gene deletion and growth assay

To study the function of *FNP1*, we generated a deletion mutant by replacing the entire gene with a hygromycin-resistance marker. Our initial effort to use homologous gene replacement strategy, which was commonly used in many of our previous Fusarium genetics studies (Fernández-Martín, et al. 2000; Shim and Woloshuk 2001; Khang, et al. 2005; Zhang, et al. 2018; Zhang, et al. 2019; Zhang, Kim, et al. 2022), was not successful, and therefore we devised a modified CRISPR/Cas9 methodology for this Fov4 gene mutation (Wang, et al. 2018). The gene-deletion mutants were confirmed by two sets of screening PCR to confirm that the *FNP1* gene was replaced with *HPH* gene (Fig. S3A). We also tested that only a single copy of the *HPH* gene was inserted into the Fov4 genome using a droplet digital (dd) PCR (Fig. S3B). These results confirmed that that the mutant strain (*Δfnp1*) had a single-copy *HPH* marker inserted into the target *FNP1* locus with no additional ectopic insertion event. The WT *FNP1* gene, driven by its native promoter, was co-transformed with a geneticin resistance marker into Δ*fnp1* mutant protoplasts to generate the complementation strain *fnp1C*. Colonies were subsequently screened by PCR, and qPCR was performed to confirm gene expression (Fig. S4).

To investigate the functional role of *FNP1* in Fov4 vegetative growth, we cultured WT, Δ*fnp1* mutant, and *FNP1*-complementated (*fnp1C*) strains on PDA, YEPD, cotton root-modified plates, and cotton root-only plates (Fig. 1). Growth diameters were measured on the 4th and 7th days for PDA, YEPD, and cotton root-modified plates, with images taken on the 7th day. No significant differences in growth rate or hyphal development were observed among the WT, Δ*fnp1*, and *FNP1* strains on these plates (Fig. 1B). For the cotton root-only plates, images were taken three weeks post-inoculation, when obvious phenotypic differences were visible. The Δ*fnp1* strain exhibited little to no aerial hyphae compared to WT, whereas the *fnp1C* strain exhibited restored WT phenotype (Fig. 1A).

**Fig. 1.**
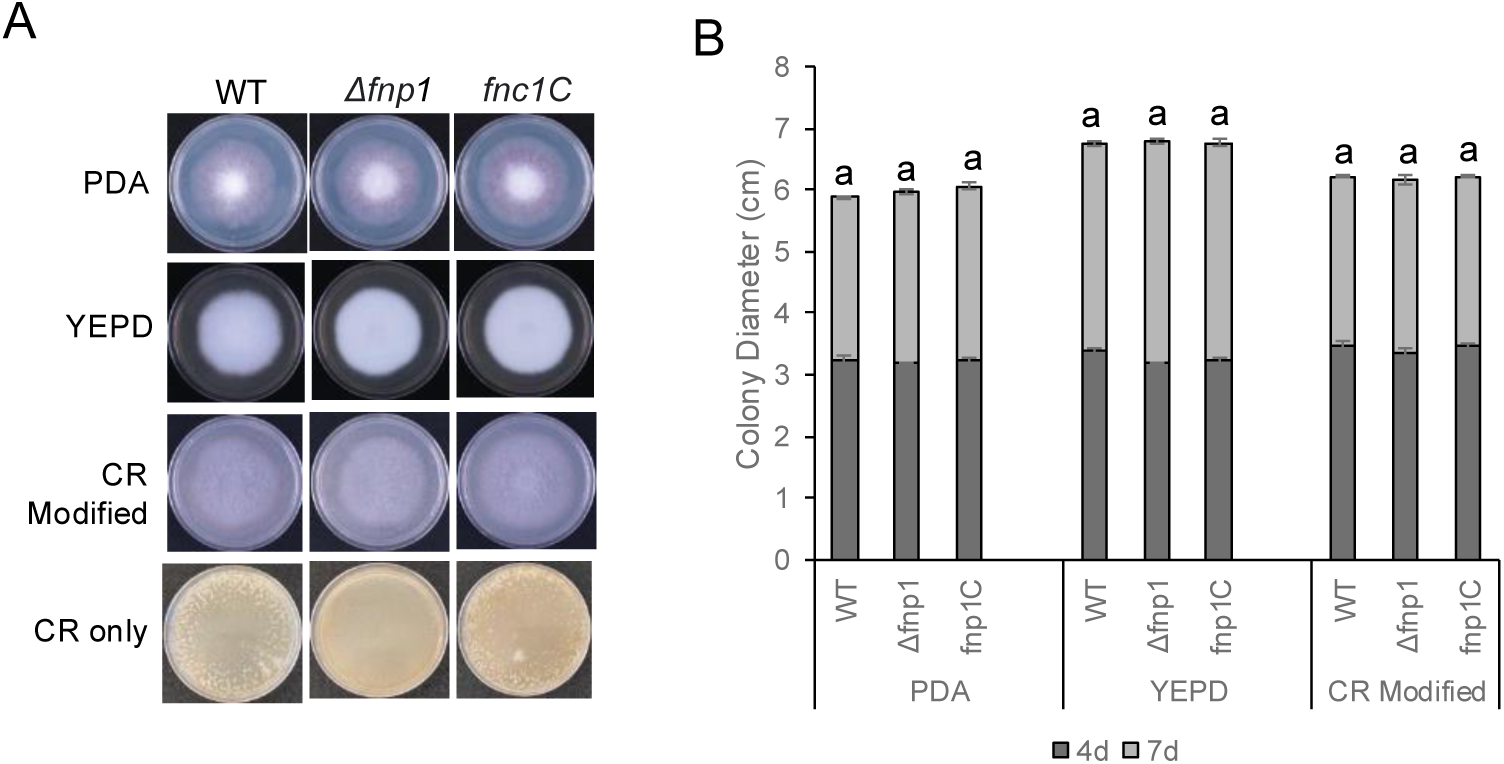
Growth assay on different media. (A) Images of 7-day-old colonies on PDA, YEPD, CR (cotton roots) Modified plates, along with 3-weeks-old colony image on CR (cotton roots) only plates. (B) Fungal colony diameter (4-day and 7-day) on PDA, YEPD, CR (cotton roots) Modified plates. Three replicates were used in this assay. Standard deviation from three independent experiments and significant differences (*P*<0.01) were analyzed by *t*-test.

Additionally, conidial production was evaluated on V8 agar plates, and no significant differences were observed among WT, Δ*fnp1*, and *fnp1C* strains (Fig. 2A). However, when we evaluated conidial germination, we observed that the Δ*fnp1* exhibited significantly slower germination compared to the WT strain (Fig. 2B). After 5 hours of incubation in 0.2× PDB medium, Δ*fnp1* strain produced significantly shorter germ tubes than the WT and *fnp1C* strains by 63.8% and 61.8%, respectively (*P* < 0.01) (Fig. 2C).

**Fig. 2.**
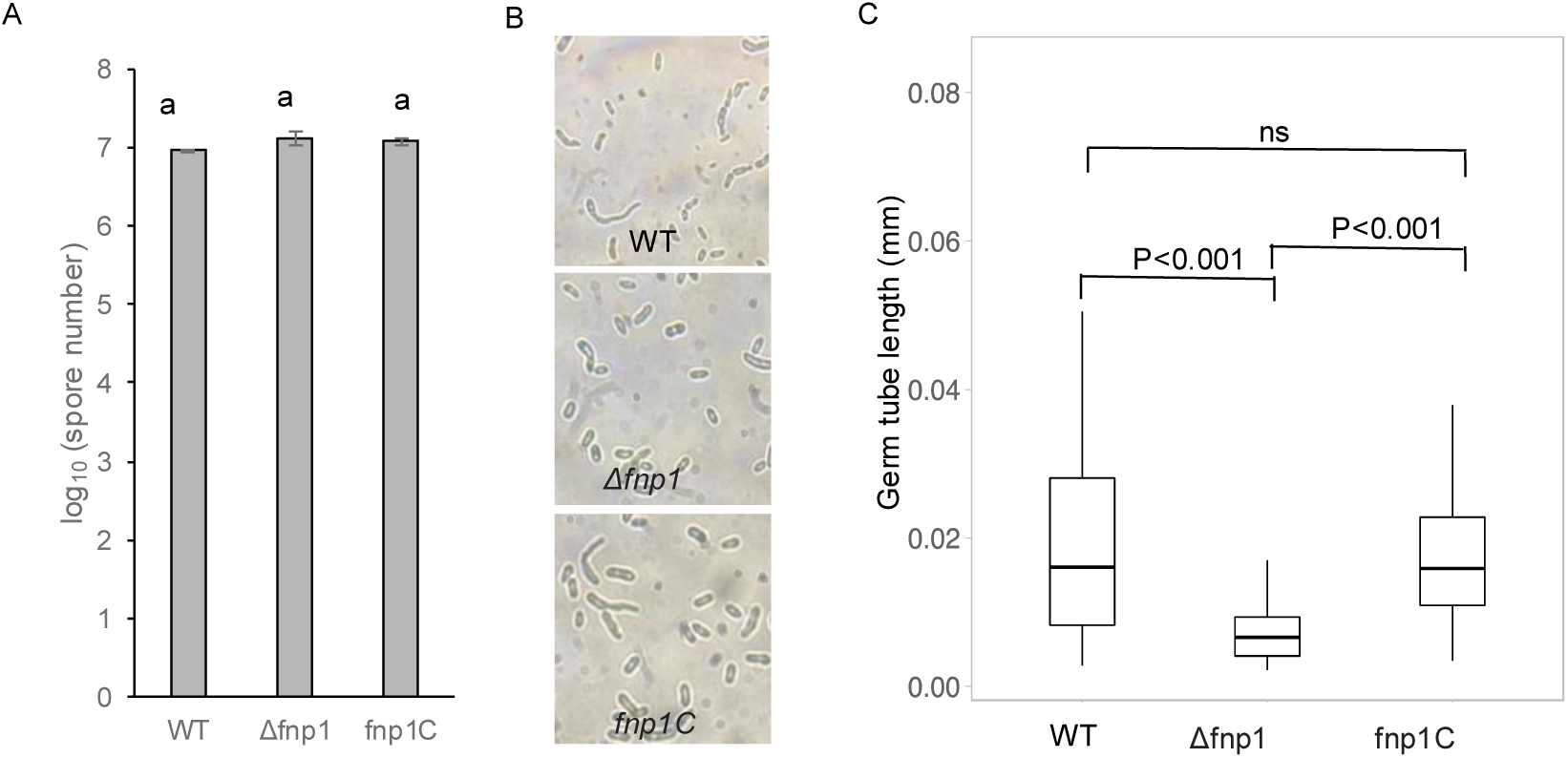
Conidial production and germination of the WT, *FNP1* gene deletion mutant (*Δfnp1*) and complementation strain (*fnp1C)*. (A) Spore production of the WT, Δ*fnp1*, and *fnp1C* strains after 10 days of growth on V8 agar plates. (B) Germinated conidia in WT, Δ*fnp1*, and *fnp1C* in 0.2× PDB were examined after 5 h of incubation with gentle shaking. (C) Conidia germinating tube length in WT, Δ*fnp1*, and *fnp1C* in 0.2× PDB were examined after 5 h of incubation with gentle shaking. Standard deviation from three independent experiments and significant differences (*P*<0.01) were analyzed by *t*-test.

### 3.3 FNP1 plays an important role in response to stressors

To investigate whether *FNP1* is involved in the response to environmental stress agents, we assessed the vegetative growth of strains on Czapek-Dox media supplemented with KCl (osmotic stress), Congo red (cell wall stress) and SDS (cell membrane stress) (Fig. 3A). The growth rate of Δ*fnp1* strain was significantly inhibited under KCl and Congo red stress conditions compared to the WT strain (Fig. 3B). The increased sensitivity to cell wall and osmotic stress, indicates that the product of *FNP1* plays a role in protecting the fungus from environmental stress.

**Fig. 3.**
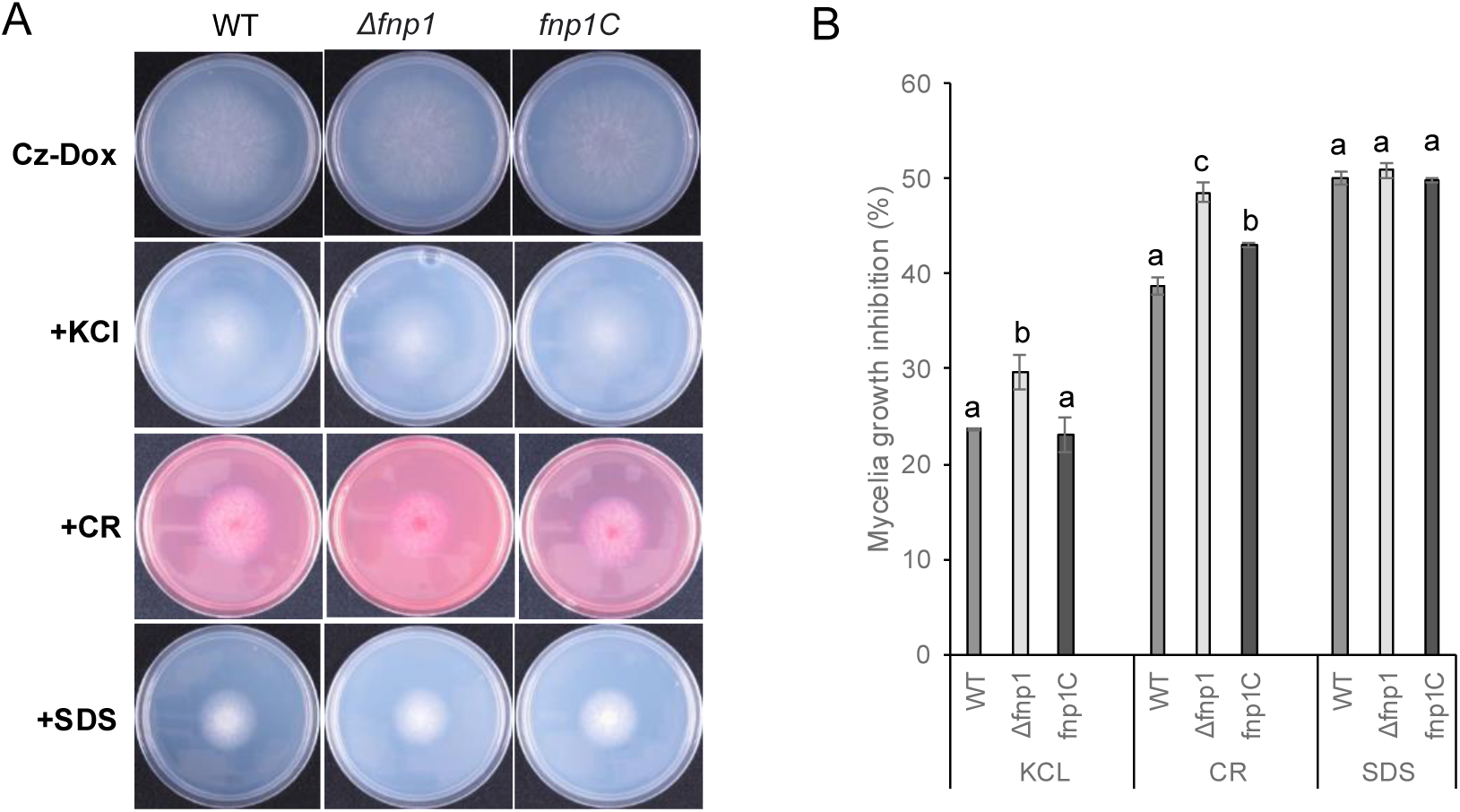
Defects of the *Δfnp1* deletion mutant in response to various stressors. (A) Five microlitres of 1×10^5^ ml^−1^ spore suspensions of WT, mutant and complemented strains were cultured on Czapek-Dox (Cz-Dox) agar amended with KCl, Congo Red and SDS for 7 days at 22°C. (B) Inhibition rate of strains grown on the media containing different stressors. Three replicates were used in this assay. The growth inhibition rate (%) was measured by (diameter of growth on Czapek-Dox agar - designated stressors)/diameter of growth on Czapek-Dox agar / diameter of growth on Czapek-Dox agar × 100. Standard deviation from three independent experiments and statistically significant differences (*P*<0.01) were analyzed by *t*-test.

### 3.4 FNP1 is important for cotton root rot virulence

To investigate whether *FNP1* plays a role in Fov4 virulence, ono-week-old cotton seedlings were inoculated with spores from the WT, Δ*fnp1*, and *fnp1C* strains using the root dip-inoculation method (Fig. 4). The disease severity was assessed 4-weeks post-inoculation. The *Δfnp1* mutant displayed a significantly lower root rot percentage and reduced root surface area compared to the WT strain. Additionally, fungal hyphae recovery was significantly higher in the WT and *fnp1C* strains than in the mutant. These data demonstrated that *FNP1* is important for cotton root rot virulence.

**Fig. 4.**
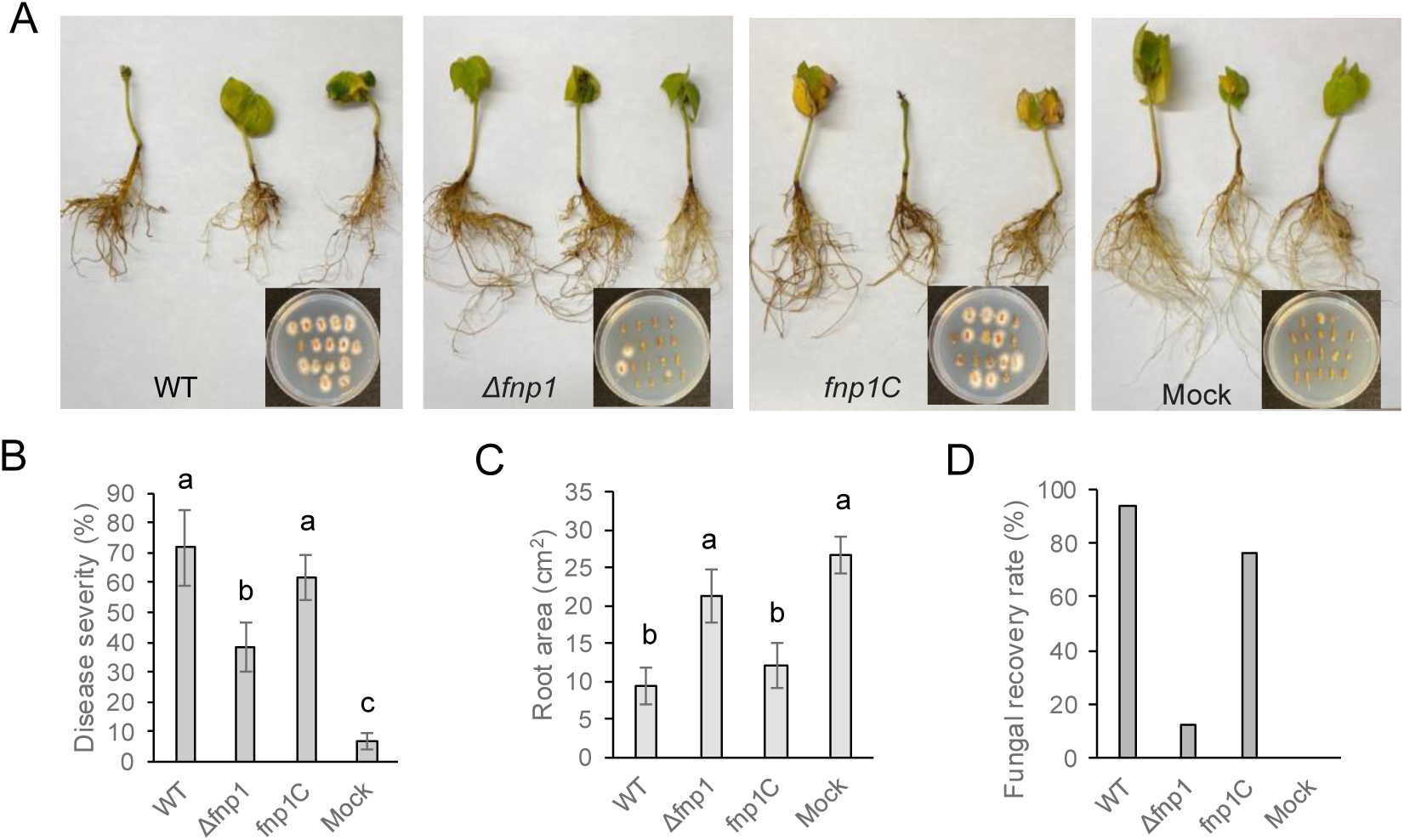
Role of *FNP1* in cotton root rot virulence. WT, Δ*fnp1* and *fnp1C* spore suspensions were inoculated on 1-week-old upland cotton roots using the root dipping method. (A) Symptoms were observed after 28-days-post inoculation. (B) Disease severity was assessed using a visual scoring scale ranging from 0 to 100% based on root discoloration. (C) Root surface area was measured using ImageJ. (D) Fungal recovery and pathogen quantification involve sterilizing root and stem segments (∼1 cm) and incubating them on PDA/KAN/AMP plates. Standard deviation from three independent experiments and statistically significant differences (*P*<0.01) were analyzed by *t*-test.

### 3.5 FNP1 is important for fusaric acid production

We tested FA production in Fov4 strains in Czapek-Dox liquid medium after a 2-week incubation. The results showed that Δ*fnp1* mutant produced dramatically lower levels of FA than the WT strain (Fig. 5A). For comparison, Beauvericin, another secondary metabolite produced by Fov4, did not show significant different production among different strains (Fig. 5A). To further understand how *FNP1* impacts FA production at the molecular level, we identified key genes within the FUB gene cluster, including *FUB1* (a polyketide synthase) and *FUB11* (a transporter involved in FA export) (Studt, et al. 2016; Hoogendoorn, et al. 2018). To examine the expression of these genes, a qPCR assay was performed on WT, Δ*fnp1*, and *fnp1C* strains cultured in cotton root liquid media for two weeks. RNA extraction followed by qPCR analysis revealed significantly reduced expression levels of *FUB1* and *FUB11* in the Δ*fnp1* strain compared to WT and *fnp1C* strains (Fig. 5B). These data demonstrated that *FNP1* influences FA production through regulation of these FUB cluster genes.

**Fig. 5.**
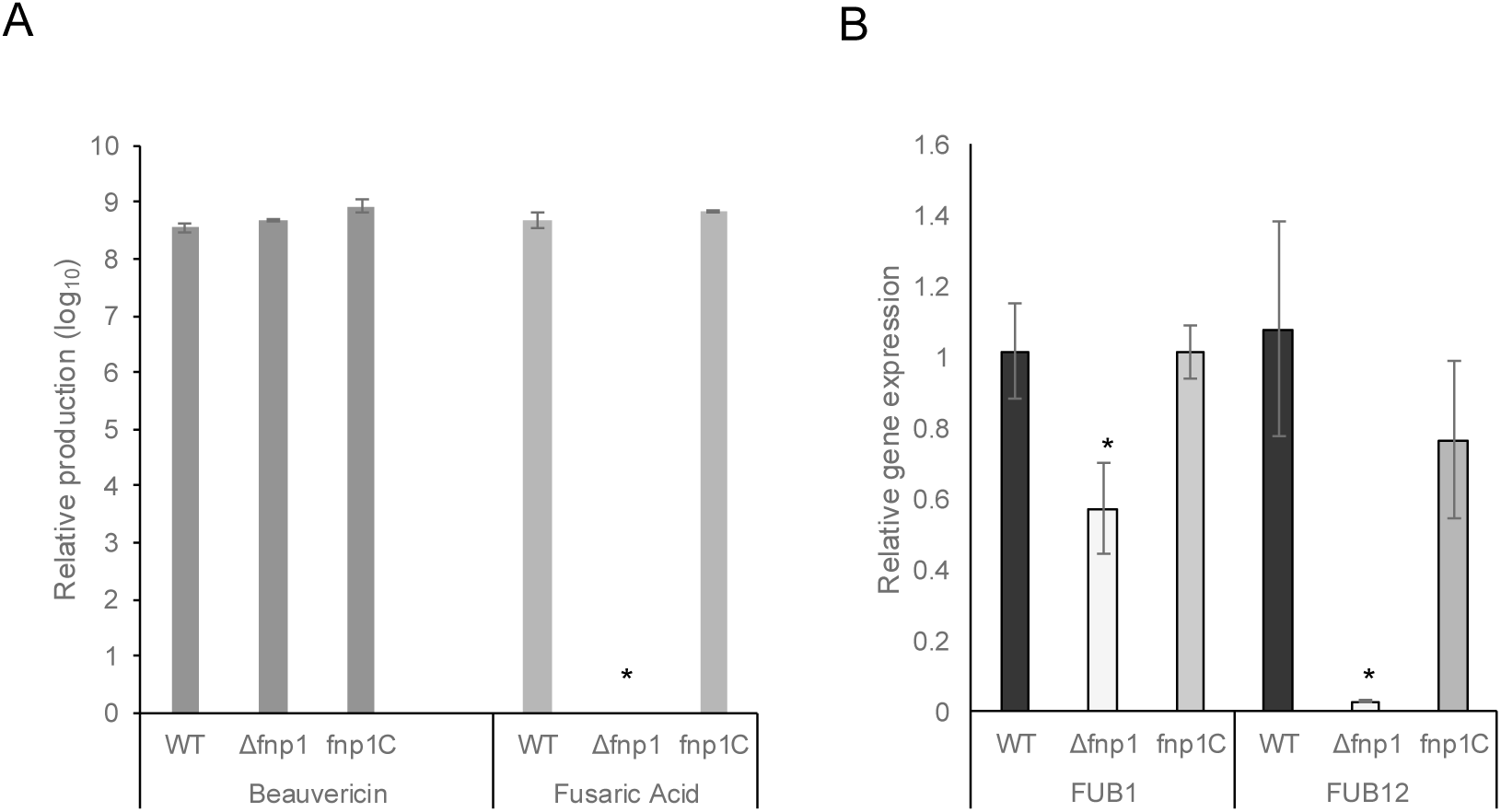
Quantification of Fusaric acid and Beauverican production in WT, *Δfnp1 and fnp1C* strains. (A) Spores of the three strains were inoculated into Czapek-Dox media incubated for 14 days. Three replicates were performed for each strain. (B) qPCR analysis of *FUB1* and *FUB12* expression levels in WT, *Δfnp1* and *fnp1C* strains. All values represent the means of three biological replications with the standard errors shown as error bars. Standard deviation from three independent experiments and statistically significant differences (*P*<0.01) were analyzed by *t*-test.

### 3.6 Infection on cotton roots using GFP tagged strain

To further confirm the impact of *FNP1* on root colonization, GFP was fused to the N-terminus of the tubulin gene under the native promoter. This construct was introduced into WT and Δ*fnp1* mutant strains, generating tubulin-GFP-WT and tubulin-GFP-Δ*fnp1* strains. Cotton roots were infected with spores from these two strains using the root dipping method. The cellular localization of the fusion proteins in cotton roots was examined 3-days post inoculation (3 dpi). Infected cotton roots were sectioned both crosswise and lengthwise at a thickness of 125 µm and analyzed by confocal microscopy. In the tubulin-GFP-WT strain, GFP signals were detected within root cortical tissue and extended into deep layers, indicating extensive intracellular hyphal colonization. In contrast, the tubulin-GFP-Δ*fnp1* strain exhibited limited hyphal progression, with fungal structures largely restricted to outer root layers. This difference was consistent across both cross-sectional and longitudinal views. Additionally, Propidium iodide (PI) staining confirmed that while WT hyphae spread through internal root tissues, Δ*fnp1* hyphae mostly localized near the root surface area (Fig.6). These observations demonstrate that *FNP1* is required for efficient colonization and deeper tissue invasion in cotton roots.

**Fig. 6.**
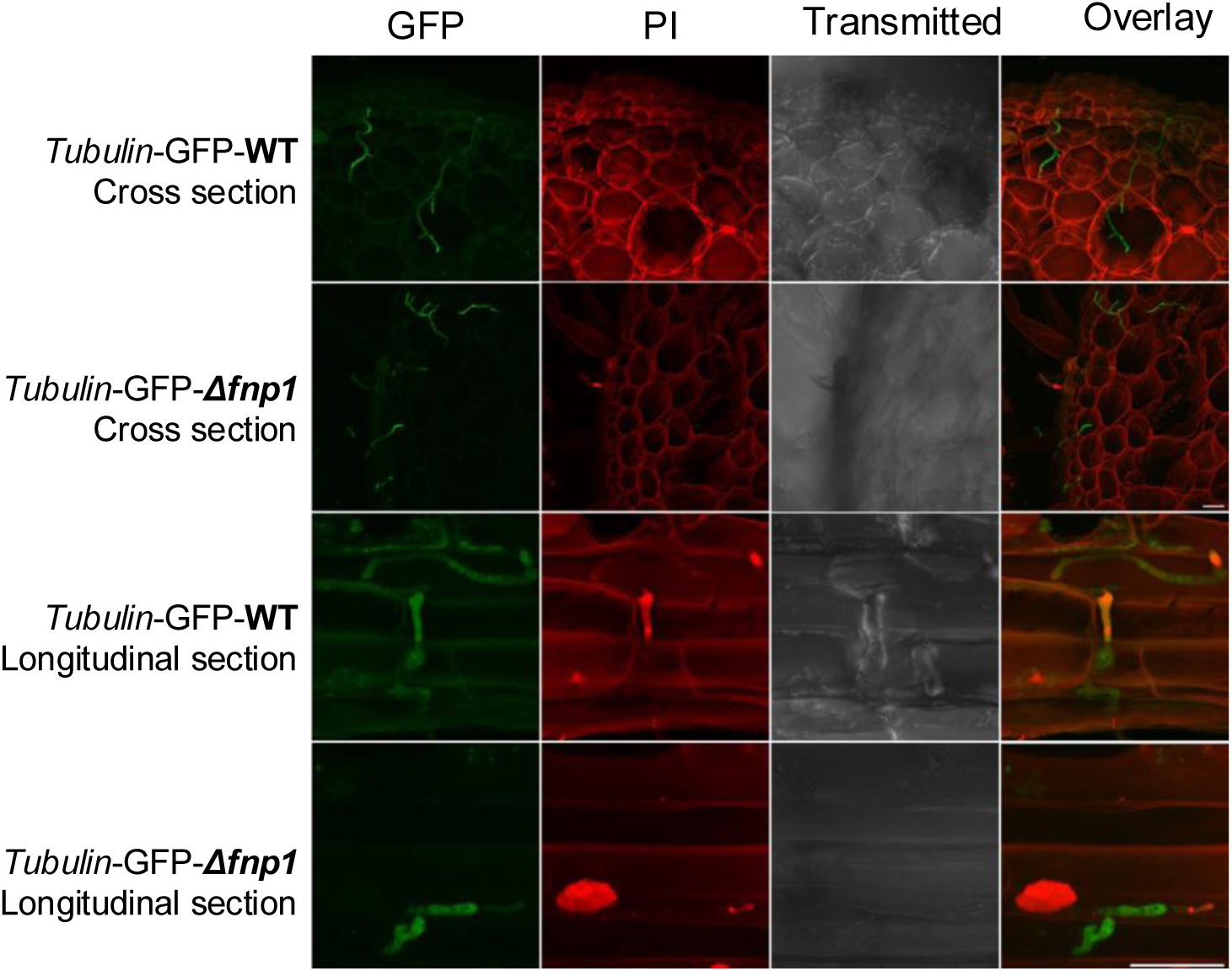
**Infection of cotton roots by GFP-tagged strains *tubulin*-GFP-WT and *tubulin*-GFP-*Δfnp1.*** Cellular localization of the fusion proteins was examined at 3-days-post inoculation (3 dpi). Cotton roots were sectioned vertically or horizontally at a thickness of 125 µm. Sections were carefully separated from agarose under a dissecting microscope and stained with propidium iodide (50 µg/mL). Imaging was taken and subsequently analyzed. Scale bars are included in the images. Scale bar = 20µm.

## 4 Discussion

The rising prevalence and geographic spread of Fov4 poses a growing threat to global cotton production. Fov4 is capable of infecting both Upland and Pima cotton varieties, with documented cases in in El Paso and Hudspeth counties in Texas and in New Mexico (Halpern, et al. 2018; Zhu, et al. 2020; Wagner, et al. 2022). Unlike other Fov races, Fov4 can infect cotton roots without the presence of RKN (Bell, et al. 2017). Furthermore, Fusarium wilt by Fov4 is considered an early season disease as the pathogen infects cotton during the seed germination and cotyledon stages, leading to root rot, seedling wilt (damping off), and death immediately after emergence (Bennett, et al. 2011; Bell, et al. 2019). In heavily infested fields, Fov4 has caused up to 90–100% seedling mortality before midseason (Zhu et al., 2021). Fov4 produces thick-walled survival structures called chlamydospores, which can persist in soil or plant residue for long periods without a host and are highly resistant to fungicides (Ulloa, et al. 2020a). Due to the limited effectiveness of crop rotation and fungicides, developing resistant cotton varieties is recognized as the most promising management strategy. Since the emergence of Fov4 in the US, breeding programs have screened thousands of lines for resistance or tolerance and several Pima cotton lines have been suggested as exhibiting resistance to Fov4 (Ulloa, et al. 2020b; Schultz 2023). However, it is important to note that Upland cotton (*Gossypium hirsutum*) constitutes approximately 97% of cotton production in the US, with Texas as the leading producer that contributes over 40% of the nation’s total output (USDA-ERS 2022). The economic significance of Upland cotton is substantial, as it serves as a vital raw material for the textile and clothing industries, significantly supporting the US agricultural economy. Achieving comparable levels of resistance in Upland cotton remains a significant challenge, there is no Upland germplasm or commercial cultivar showing Fov4 resistant/tolerant available in the US (Ulloa, et al. 2020a).

In this study, our aim was to investigate genetic factors that enable Fov4 to be such an aggressive Fov race on cotton seedlings. Recognizing that secondary metabolites play crucial roles in determining the virulence in fungal pathogens, we performed a comparative analysis of Fov1 and Fov4 genomes to identify a unique NRPS gene, *FNP1*, in Fov4. A CRISPR/Cas9 system was employed to generate gene deletion mutants and GFP labeling. Growth assays showed that the *FNP1* deletion mutant exhibited slower conidia germination and reduced growth under stress conditions. Additionally, the mutant demonstrated significantly decreased virulence in causing cotton root rot, accompanied by a notable reduction in fusaric acid production, highlighting the essential role of *FNP1* in Fov4 pathogenicity. Fusaric acid is a mycotoxin produced by various *Fusarium* species, particularly those species known to be plant pathogens (Rani, et al. 2009; Singh, et al. 2017). It is generally considered to have moderate levels of toxicity on plants but can act synergistically with other toxic secondary metabolites leading to cell death and necrosis.

What was perplexing about our outcome was that the putative *FNP1* secondary metabolite serving as a regulator or stimulant for fusaric acid production in Fov4. It is intriguing to hypothesize that NRPS and PKS pathways are cross-talking to regulate virulence mechanisms in a fungal pathogen.

Previous studies identified several polyketides produced by *Fusarium oxysporum*, including the nonaketide naphthazarin quinones, bikaverin, and norbikaverin (Bell, et al. 2003). However, these compounds play only limited roles in virulence. Among the secondary metabolites produced by Fov4, FA has been detected in culture filtrates, and it is reported to play a crucial role in symptom development in *Fusarium oxysporum* f. sp. *lycopersici* (Fol) (Mapuranga, et al. 2022; Zhu, et al. 2022; Zhu, et al. 2023). Additionally, *Fusarium oxysporum* has been reported to produce other mycotoxins, such as beauvericin, enniatins A and zearalenone (ZEA) (Koenning, et al. 2004; Song, et al. 2008; Beev, et al. 2013). To further investigate the contribution of these secondary metabolites to Fov4 virulence, we assessed the presence and abundance of FA, beauvericin, ENN A, and ZEA in culture filtrates. Our analysis indicated that ENN A and ZEA were either absent or detected only in trace amounts (data not shown), suggesting they are unlikely to contribute significantly to Fov4 virulence. Therefore, we focused on FA and beauvericin by comparing their production between WT and *FNP1* deletion mutant strains to gain deeper insights into the pathogenic mechanisms of Fov4. Our results revealed that the *FNP1* deletion mutant exhibited significantly reduced levels of FA compared to the WT strain, indicating that *FNP1* plays an important role in FA biosynthesis. In contrast, no significant differences in beauvericin production were observed between WT and mutant strains, suggesting that *FNP1* is not involved in the beauvericin biosynthesis pathway. These findings imply that *FNP1* is functionally or transcriptionally linked to the FA biosynthetic pathway but does not participate in BEA biosynthesis. Secondary metabolite biosynthesis in filamentous fungi is typically organized into discrete BGCs, each regulated by specific transcription factors, allowing for pathway-specific control. This functional and genetic independence has been demonstrated in several fungal species. For example, in *F. graminearum*, deletion of the histidine kinase *FgOs1*, part of the HOG pathway, led to a dramatic decrease in the red pigment aurofusarin, while trichothecene production was not impacted (Ochiai, et al. 2007). The authors proposed that additional downstream elements, other than those in the osmoregulatory MAPK pathway linked to the upstream phosphorelay system, may specifically regulate pigment biosynthesis without affecting trichothecene production. Similarly, in *Acremonium chrysogenum*, disruption of the *CPCR1*gene led to reduced levels of penicillin N. However, cephalosporin levels remained unchanged, indicating that *CPCR1* does not regulate the late-stage genes in the cephalosporin biosynthetic pathway (Schmitt, et al. 2004). These studies illustrate that secondary metabolic pathways in filamentous fungi are regulated independently, providing a mechanistic basis for selectively targeting specific metabolites involved in virulence or environmental adaptation without disrupting other secondary metabolic processes.

Our findings indicate that *FNP1* contributes to fungal protection against environmental stress, as evidenced by the increased sensitivity of the *Δfnp1* mutant to both cell wall stress (Congo red) and osmotic stress (KCl) (Fig. 3). This increased stress sensitivity, combined with the reduced virulence of the mutant, suggests a potential link between stress tolerance and fungal pathogenicity. Secondary metabolites are well known to play diverse roles in fungal biology, including providing structural support, regulating development, and defending against environmental challenges. The product of *FNP1* likely contributes to one or more of these protective functions enabling the fungus to cope with stress encountered during infection or environmental exposure. Many studies have demonstrated the critical roles of fungal secondary metabolites, particularly those synthesized by non-ribosomal peptide synthetases (NRPSs), in stress adaptation and pathogenicity (Panaccione, et al. 1992; Haese, et al. 1993). For example, in *Cochliobolus heterostrophus*, deletion of *NPS6*, which is responsible for the biosynthesis of a siderophore-like compound, results in increased sensitivity to oxidative stress. The *nps6* mutant showed significantly reduced growth in the presence of hydrogen peroxide, highlighting its essential role in oxidative stress resistance and virulence (Lee, et al. 2005). Similarly, in *Alternaria brassicicola*, loss of *AbNPS2* leads to conidia with abnormal morphology, including inflated cell walls and elevated lipid body accumulation. These mutants also displayed increased susceptibility to UV-induced damage, with a 25% reduction in germination following UV exposure compared to the WT, indicating a broader role for NRPS-derived metabolites in protecting against both abiotic and biotic stressors (KIM, et al. 2007). Together, these studies, along with our findings, support the view that NRPS-derived secondary metabolites are integral to fungal adaptation to a variety of environmental and biological stresses, and that this stress resilience may be closely tied to virulence.

As soil-borne pathogens interact with specific microbial taxa in the soil environment, root microbiomes play a crucial role in plant health and disease resistance as reported in important crops such as rice, maize, crops and tomatoes (Sharma; Meshram and Adhikari 2024). The composition of the rhizosphere microbiota (*i.e.* soil associated directly with plant roots) plays more important role in modulating plant defense responses, potentially enhancing or suppressing the plant resistance against pathogen infection (Doornbos, et al. 2012; de Faria, et al. 2021). It has been reported that rhizosphere microorganisms act as antagonists against pathogenic *F. oxysporum* on *Crocus sativus* (Zhang, Lu, et al. 2022). A study also revealed that Fov directly and consistently altered the rhizosphere microbiome, but the biocontrol agents enabled microbial assemblages to resist pathogenic stress on cotton (Qiu, et al. 2022). It is conceivable that fungal secondary metabolites play an important, but yet-to-be determined, role in the soil microbiome. Notably, we can propose that soilborne pathogenic fungi are potent ecosystem modulators that can influence the overall microbial functioning. And the effect of fungi beyond the rhizosphere to influence the overall soil functions is seldom understood. Understanding these complex interactions is crucial for developing integrated disease management strategies that manipulate the soil microbiome to mitigate the impact of Fov4 on cotton production. Studies have shown that soils suppressive to Fusarium diseases harbor specific microbial communities that inhibit pathogen growth while promoting plant health. Whether fungal secondary metabolites play direct or indirect role is not known. Further research is required to explore the role of soil microbiome dynamics in effectively managing Fusarium wilt in cotton.

## Supplementary Materials

The following supporting information can be downloaded online. Fig. S1. Identification of Fov4-specific secondary metabolite biosynthetic genes. Secondary metabolite biosynthetic gene clusters in Fov4 (column 1) and Fov1 (column 2) were predicted using antiSMASH. BLASTp analysis was used to identify genes unique to Fov4. A nonribosomal peptide synthetase cluster in Fov4 (NRPS12.2, highlighted in red) was predicted by antiSMASH and identified as *Fov4* specific. Fig. S2. Domain architecture of the Fnp1 protein. The predicted 2252-amino-acid protein contains three conserved condensation (C) domains, one adenylation (A) domain, and two phosphopantetheine (Krappmann) binding domains, as identified by antiSMASH analysis. Fig. S3. Screening and confirmation of *FNP1* knockout mutant using PCR and ddPCR.

(A) Initial screening of the *FNP1* knockout mutant was performed using ORF-screen/F and ORF-screen/R primers to assess the presence or absence of the *FNP1* gene. (B) Insertion of the *HPH* (hygromycin resistance) gene was confirmed using the upstream LF/F/screen and HYG/F primers. (C) Droplet digital PCR (ddPCR) was used to verify a single insertion of the *HPH* gene in the selected mutant (Marked with asterisk). Fov4 *b*-*tubulin* gene served as a control. Fig. S4. Generation and confirmation of *FNP1* complementation strain. (A) The *FNP1* gene, under the control of its native promoter, was co-transformed with a geneticin resistance gene into mutant protoplasts. Drug-resistant colonies were screened by PCR to verify the presence of the *FNP1* gene. (B) Expression of *FNP1* in the selected complementation candidate (Marked with red asterisk) was confirmed by quantitative PCR (qPCR).

## Acknowledgements

This research was supported by the United States Department of Agriculture (2022-67013-36836) and National Institute of Food and Agriculture (2023-67013-40174). Authors declare no conflict of interest.

## Author Contributions

Huan Zhang., Yi Zhou., and Won Bo Shim conceived and designed the experiments. Huan Zhang, Yi Zhou and Caleb Oliver Bedsole performed experiments and analyses. Huan Zhang, Yi Zhou, Caleb Oliver Bedsole and Won Bo Shim wrote the manuscript.

---

**Fig. S1.**
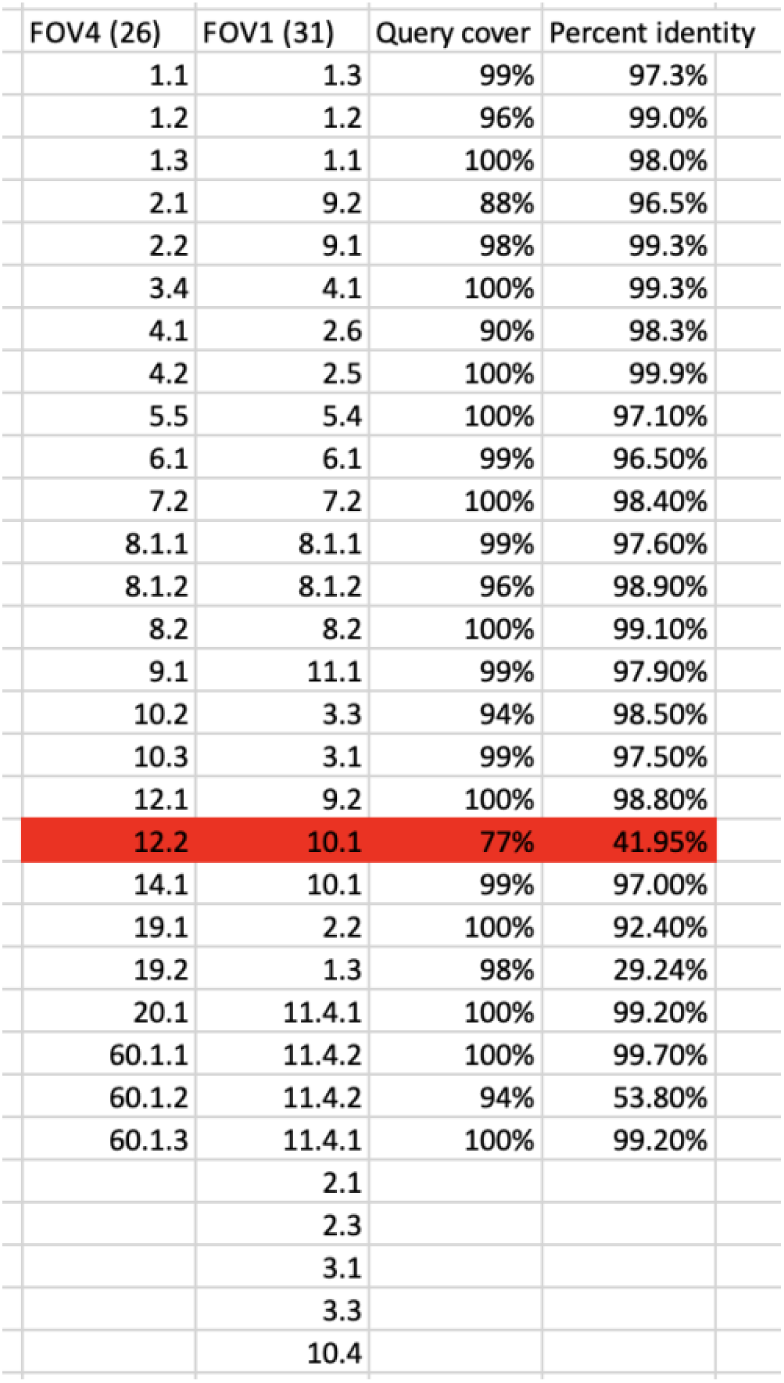
Identification of Fov4-specific secondary metabolite biosynthetic genes. Secondary metabolite biosynthetic gene clusters in Fov4 (column 1) and Fov1 (column 2) were predicted using antiSMASH. BLASTp analysis was used to identify genes unique to Fov4. A nonribosomal peptide synthetase cluster in Fov4 (NRPS12.2, highlighted in red) was predicted by antiSMASH and identified as Fov4 specific.

**Fig. S2.**
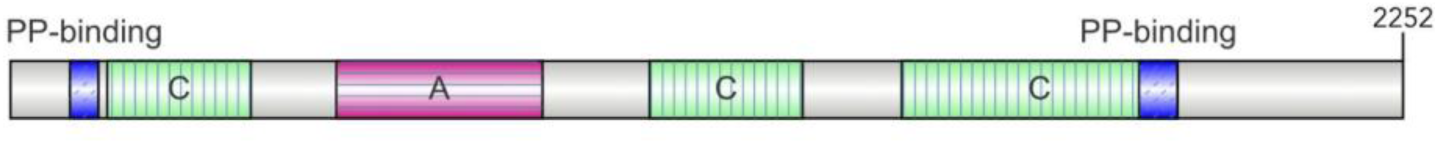
Domain architecture of the Fnp1 protein. The predicted 2252-amino-acid protein contains three conserved condensation (C) domains, one adenylation (A) domain, and two phosphopantetheine (PP) binding domains, as identified by antiSMASH analysis.

**Fig. S3.**
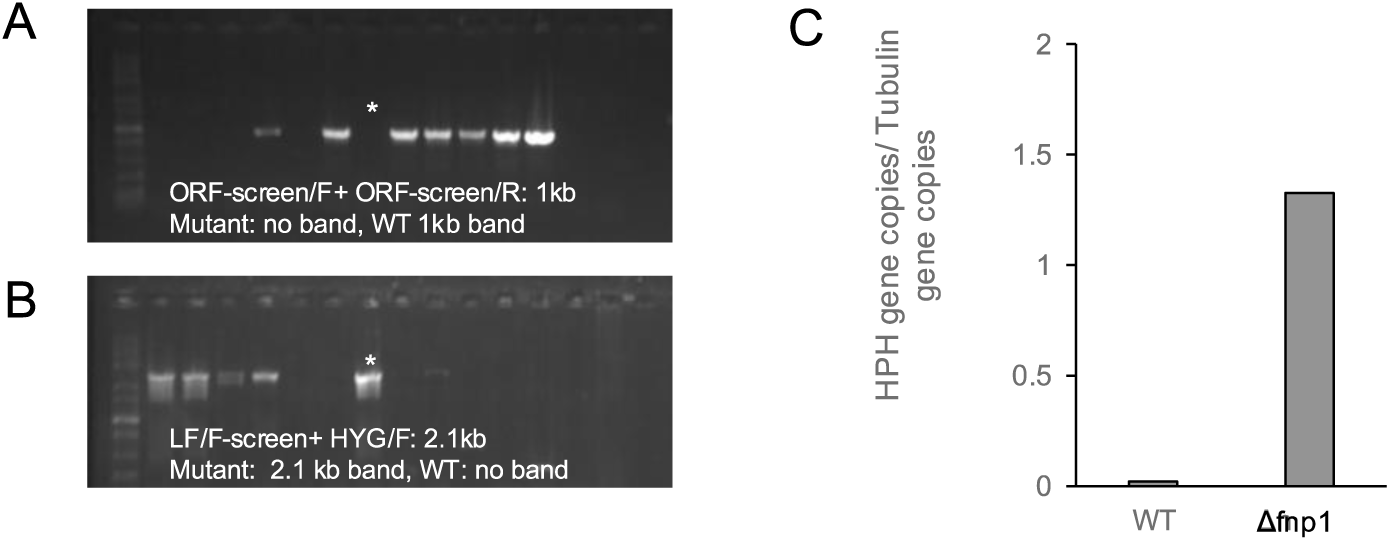
Screening and confirmation of *FNP1* knockout mutant using PCR and ddPCR. (A) Initial screening of the *FNP1* knockout mutant was performed using ORF-screen/F and ORF-screen/R primers to assess the presence or absence of the *FNP1* gene. (8) Insertion of the *HPH* (hygromycin resistance) gene was confirmed using the upstream LF/F/screen and HYG/F primers. (C) Droplet digital PCR (ddPCR) was used to verify a single insertion of the *HPH* gene in the selected mutant (Marked with asterisk). Fov4 *b-tubulin* gene served as a control.

**Fig. S4.**
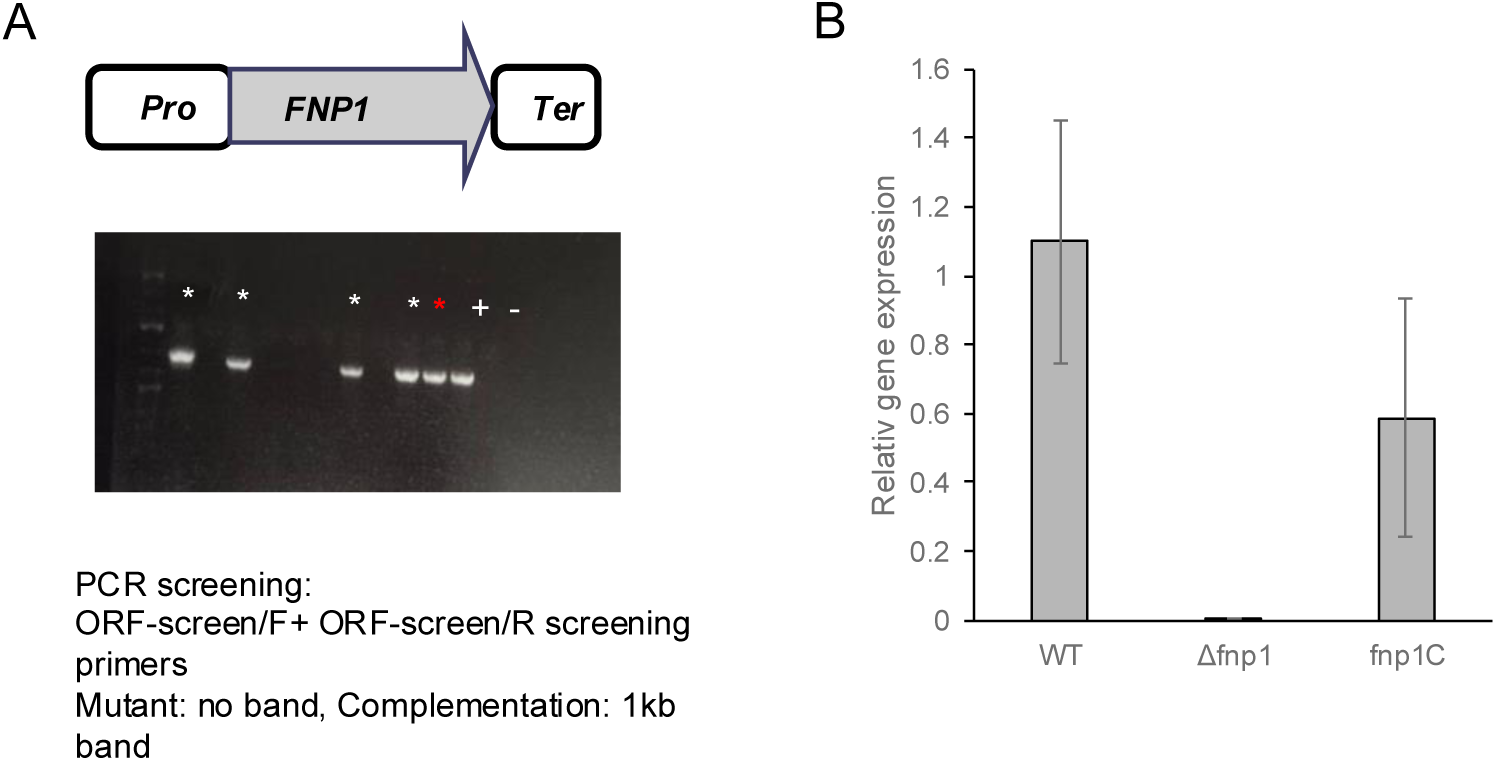
Generation and confirmation of *FNP1* complementation strain. (A) The *FNP1* gene, under the control of its native promoter, was co-transformed with a geneticin resistance gene into mutant protoplasts. Drug-resistant colonies were screened by PCR to verify the presence of the *FNP1* gene. (B) Expression of *FNP1* in the selected complementation candidate (Marked with red asterisk) was confirmed by quantitative PCR (qPCR).

**Table S1:**
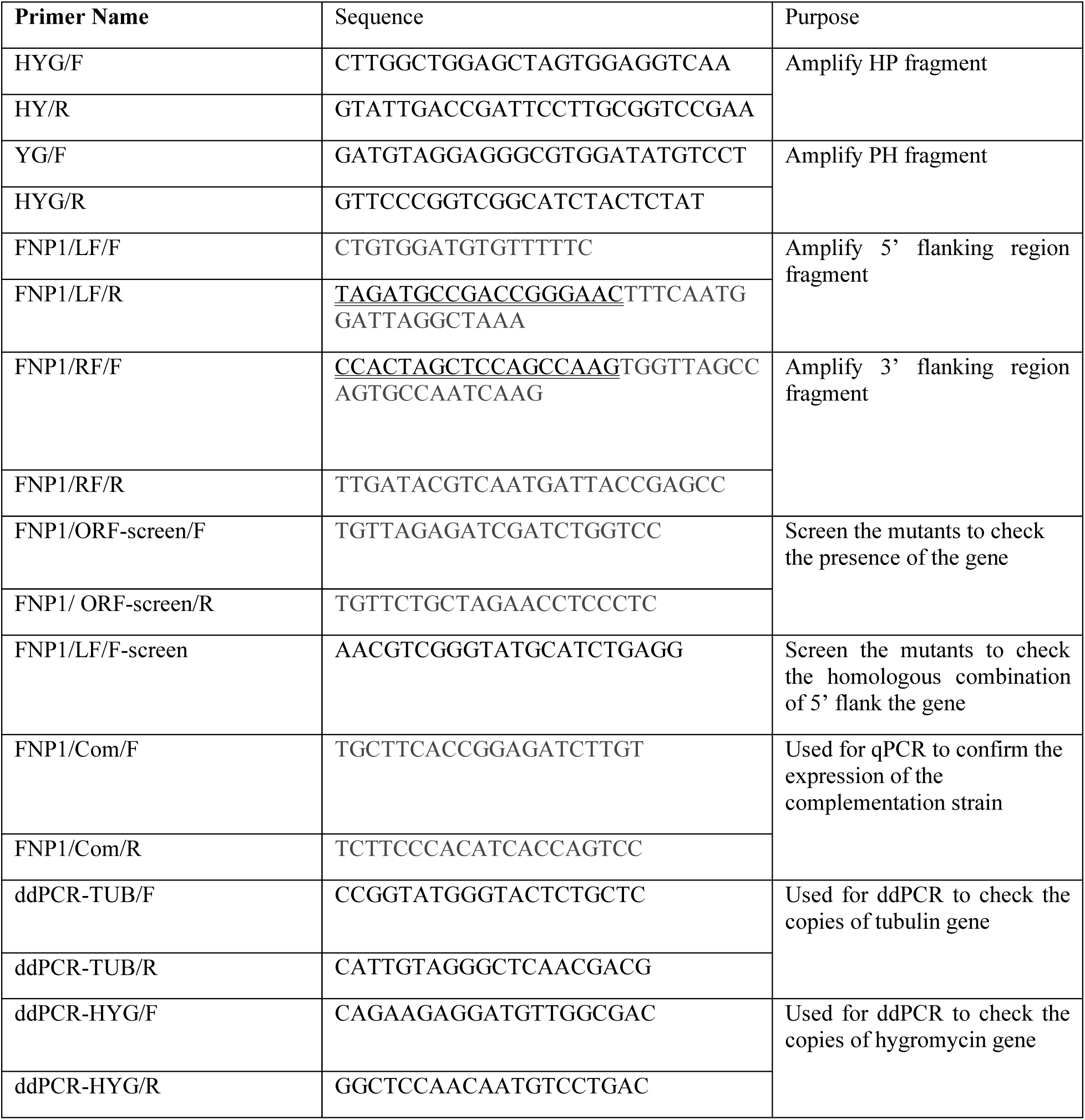
Primers used in this study.

## Notes

### Competing Interest Statement

The authors have declared no competing interest.

## REFERENCES

Armstrong GM, Armstrong JK. 1960. American, Egyptian, and Indian cotton-wilt fusaria: their pathogenicity and relationship to other wilt fusaria: US Department of Agriculture.

Ayubov MS, Abdurakhmonov IY, Makamov AK, Mamajonov BO, Yusupov AN, Obidov NS, Bashirxonov ZH, Murodov AA, Darmanov MM, Ubaydullaeva KA. 2024. Recent Studies on Fusarium Wilt in Cotton. Fusarium-Recent Studies.

Bacon C, Porter J, Norred W, Leslie J. 1996. Production of fusaric acid by Fusarium species. Applied and Environmental Microbiology 62:4039–4043.

Beev G, Denev S, Bakalova D. 2013. Zearalenone-producing activity of *Fusarium graminearum* and *Fusarium oxysporum* isolated from Bulgarian wheat. Bulgarian Journal of Agricultural Science 19:255–259.

Bell AA, Gu A, Olvey J, Wagner TA, Tashpulatov JJ, Prom S, Quintana J, Nichols RL, Liu J. 2019. Detection and characterization of *Fusarium oxysporum* f. sp. vasinfectum VCG0114 (Race 4) isolates of diverse geographic origins. Plant Disease 103:1998–2009.

Bell AA, Kemerait RC, Ortiz CS, Prom S, Quintana J, Nichols RL, Liu J. 2017. Genetic diversity, virulence, and Meloidogyne incognita interactions of *Fusarium oxysporum* isolates causing cotton wilt in Georgia. Plant Disease 101:948–956.

Bell AA, Wheeler MH, Liu J, Stipanovic RD, Puckhaber LS, Orta H. 2003. United States Department of Agriculture—Agricultural Research Service studies on polyketide toxins of *Fusarium oxysporum* f. sp. *vasinfectum*: potential targets for disease control. Pest Management Science: Formerly Pesticide Science 59:736–747.

Bennett R, Spurgeon D, DeTar W, Gerik J, Hutmacher R, Hanson B. 2011. Efficacy of four soil treatments against *Fusarium oxysporum* f. sp. *vasinfectum* race 4 on cotton. Plant Disease 95:967–976.

Blin K, Shaw S, Kloosterman AM, Charlop-Powers Z, Van Wezel GP, Medema MH, Weber T. 2021. antiSMASH 6.0: improving cluster detection and comparison capabilities. Nucleic acids research 49:W29–W35.

Cai Y-M, Dudley QM, Patron NJ. 2021. Measurement of transgene copy number in plants using droplet digital PCR. Bio-protocol 11:e4075–e4075.

Chaudhry MR, Guitchounts A. 2003. Cotton facts: International Cotton Advisory Committee Washington, DC.

Cianchetta AN, Allen TW, Hutmacher RB, Kemerait RC, Kirkpatrick TL, Lawrence GW, Lawrence KS, Mueller JD, Nichols RL, Olsen MW. 2015. Survey of *Fusarium oxysporum* f. sp. vasinfectum in the United States.

Cianchetta AN, Davis R. 2015. Fusarium wilt of cotton: Management strategies. Crop Protection 73:40–44.

Colyer P, Kirkpatrick T, Caldwell W, Vernon P. 1997. Influence of nematicide application on the severity of the root-knot nematode-Fusarium wilt disease complex in cotton. Plant Disease 81:66–70.

Constable G, Bange M. 2015. The yield potential of cotton (*Gossypium hirsutum* L.). Field Crops Research 182:98–106.

Crutcher FK, Liu J, Puckhaber LS, Stipanovic RD, Bell AA, Nichols RL. 2015. FUBT, a putative MFS transporter, promotes secretion of fusaric acid in the cotton pathogen *Fusarium oxysporum* f. sp. *vasinfectum*. Microbiology 161:875–883.

Crutcher FK, Puckhaber LS, Stipanovic RD, Bell AA, Nichols RL, Lawrence KS, Liu J. 2017. Microbial resistance mechanisms to the antibiotic and phytotoxin fusaric acid. Journal of chemical ecology 43:996–1006.

Davis R, Colyer P, Rothrock C, Kochman J. 2006. Fusarium wilt of cotton: population diversity and implications for management. Plant Disease 90:692–703.

de Faria MR, Costa LSAS, Chiaramonte JB, Bettiol W, Mendes R. 2021. The rhizosphere microbiome: functions, dynamics, and role in plant protection. Tropical Plant Pathology 46:13–25.

Desjardins A, Proctor R. 2007. Molecular biology of Fusarium mycotoxins. International journal of food microbiology 119:47–50.

Doornbos RF, van Loon LC, Bakker PA. 2012. Impact of root exudates and plant defense signaling on bacterial communities in the rhizosphere. A review. Agronomy for Sustainable Development 32:227–243.

Ekwomadu TI, Akinola SA, Mwanza M. 2021. Fusarium mycotoxins, their metabolites (free, emerging, and masked), food safety concerns, and health impacts. International Journal of Environmental Research and Public Health 18:11741.

El-Zik K, Thaxton P. 1998. Integrated management of cotton wilt diseases.

Fernández-Martín R, Cerdá-Olmedo E, Avalos J. 2000. Homologous recombination and allele replacement in transformants of *Fusarium fujikuroi*. Molecular and General Genetics MGG 263:838–845.

Gaumann E. 1957. Fusaric acid as a wilt toxin.

Go EB, Kim LJ, Nelson HM, Ohashi M, Tang Y. 2021. Biosynthesis of the fusarium mycotoxin (−)-sambutoxin. Organic letters 23:7819–7823.

Haese A, Schubert M, Herrmann M, Zocher R. 1993. Molecular characterization of the enniatin synthetase gene encoding a multifunctional enzyme catalysing N-methyldepsipeptide formation in Fusarium scirpi. Molecular Microbiology 7:905–914.

Halpern H, Bell A, Wagner T, Liu J, Nichols R, Olvey J, Woodward J, Sanogo S, Jones C, Chan C. 2018. First report of Fusarium wilt of cotton caused by *Fusarium oxysporum* f. sp. *vasinfectum* race 4 in Texas, USA. Plant Disease 102:446–446.

Halpern HC, Qi P, Kemerait RC, Brewer MT. 2020. Genetic diversity and population structure of races of *Fusarium oxysporum* causing cotton wilt. G3: Genes, Genomes, Genetics 10:3261–3269.

Hoh DZ, Lee H-H, Wada N, Liu W-A, Lu MR, Lai C-K, Ke H-M, Sun P-F, Tang S-L, Chung W-H. 2022. Comparative genomic and transcriptomic analyses of trans-kingdom pathogen *Fusarium solani* species complex reveal degrees of compartmentalization. BMC biology 20:236.

Hoogendoorn K, Barra L, Waalwijk C, Dickschat JS, Van der Lee TA, Medema MH. 2018. Evolution and diversity of biosynthetic gene clusters in Fusarium. Frontiers in microbiology 9:1158.

Hussain M, Gao X, Qin D, Qin X, Wu G. 2023. Role of biotic and abiotic factors for sustainable cotton production. In. Best Crop Management and Processing Practices for Sustainable Cotton Production: IntechOpen.

Inc. A. 2025. Adobe Photoshop. San Jose, CA: Adobe Inc.

Ismaiel AA, Papenbrock J. 2015. Mycotoxins: producing fungi and mechanisms of phytotoxicity. Agriculture 5:492–537.

Khan MA, Wahid A, Ahmad M, Tahir MT, Ahmed M, Ahmad S, Hasanuzzaman M. 2020. World cotton production and consumption: An overview. Cotton production and uses: Agronomy, crop protection, and postharvest technologies:1–7.

Khang CH, Park S-Y, Lee Y-H, Kang S. 2005. A dual selection based, targeted gene replacement tool for *Magnaporthe grisea* and *Fusarium oxysporum*. Fungal Genetics and Biology 42:483–492.

Kim KH, Cho Y, La Rota M, Cramer Jr RA, Lawrence CB. 2007. Functional analysis of the *Alternaria brassicicola* non-ribosomal peptide synthetase gene *AbNPS2* reveals a role in conidial cell wall construction. Molecular plant pathology 8:23–39.

Kim Y, Hutmacher R, Davis R. 2005. Characterization of California isolates of *Fusarium oxysporum* f. sp. *vasinfectum*. Plant Disease 89:366–372.

Koenning SR, Wrather JA, Kirkpatrick TL, Walker NR, Starr JL, Mueller JD. 2004. Plant-parasitic nematodes attacking cotton in the United States: Old and emerging production challenges. Plant Disease 88:100–113.

Krappmann S. 2007. Gene targeting in filamentous fungi: the benefits of impaired repair. Fungal biology reviews 21:25–29.

Lee B-N, Kroken S, Chou DY, Robbertse B, Yoder O, Turgeon BG. 2005. Functional analysis of all nonribosomal peptide synthetases in Cochliobolus heterostrophus reveals a factor, *NPS6*, involved in virulence and resistance to oxidative stress. Eukaryotic Cell 4:545–555.

Lin C, Feng X-l, Liu Y, Li Z-c, Li X-Z, Qi J. 2023. Bioinformatic analysis of secondary metabolite biosynthetic potential in pathogenic Fusarium. Journal of Fungi 9:850.

Ma L-J, Van Der Does HC, Borkovich KA, Coleman JJ, Daboussi M-J, Di Pietro A, Dufresne M, Freitag M, Grabherr M, Henrissat B. 2010. Comparative genomics reveals mobile pathogenicity chromosomes in Fusarium. Nature 464:367–373.

Mapuranga J, Chang J, Zhang L, Zhang N, Yang W. 2022. Fungal secondary metabolites and small RNAs enhance pathogenicity during plant-fungal pathogen interactions. Journal of Fungi 9:4.

Merga W. 2018. Measuring and analysis of plant diseases. International Journal of Research Studies in Agricultural Sciences 4:1–8.

Meshram S, Adhikari TB. 2024. Microbiome-mediated strategies to manage major soil-borne diseases of tomato. Plants 13:364.

Ochiai N, Tokai T, Nishiuchi T, Takahashi-Ando N, Fujimura M, Kimura M. 2007. Involvement of the osmosensor histidine kinase and osmotic stress-activated protein kinases in the regulation of secondary metabolism in *Fusarium graminearum*. Biochemical and Biophysical Research Communications 363:639–644.

Ortiz CS, Bell AA, Magill CW, Liu J. 2017. Specific PCR detection of *Fusarium oxysporum* f. sp. *vasinfectum* California race 4 based on a unique Tfo1 insertion event in the PHO gene. Plant Disease 101:34–44.

Panaccione DG, Scott-Craig JS, Pocard J-A, Walton JD. 1992. A cyclic peptide synthetase gene required for pathogenicity of the fungus *Cochliobolus carbonum* on maize. Proceedings of the National Academy of Sciences 89:6590–6594.

Porter JK, Bacon CW, Wray EM, Hagler Jr WM. 1995. Fusaric acid in *Fusarium moniliforme* cultures, corn, and feeds toxic to livestock and the neurochemical effects in the brain and pineal gland of rats. Natural Toxins 3:91–100.

Qiu Z, Verma JP, Liu H, Wang J, Batista BD, Kaur S, de Araujo Pereira AP, Macdonald CA, Trivedi P, Weaver T. 2022. Response of the plant core microbiome to *Fusarium oxysporum* infection and identification of the pathobiome. Environmental Microbiology 24:4652–4669.

Rani T, Savitha R, Lavanya L, Kamalalochani S, Bharathiraja B. 2009. An overview of Fusaric acid production. Adv. Biotech 8:18–22.

Schmitt EK, Bunse A, Janus D, Hoff B, Friedlin E, Kürnsteiner H, Kück U. 2004. Winged helix transcription factor *CPCR1* is involved in regulation of β-lactam biosynthesis in the fungus *Acremonium chrysogenum*. Eukaryotic Cell 3:121–134.

Schneider CA, Rasband WS, Eliceiri KW. 2012. NIH Image to ImageJ: 25 years of image analysis. Nature methods 9:671–675.

Schultz A. 2023. Breeding for Host Plant Resistance to FOV4 in Cotton.

Seo S, Pokhrel A, Coleman JJ. 2020. The genome sequence of five genotypes of *Fusarium oxysporum* f. sp. *vasinfectum*: A resource for studies on Fusarium wilt of cotton. Molecular Plant-Microbe Interactions 33:138–140.

Shahrajabian MH, Sun W, Cheng Q. 2020. Considering white gold, cotton, for its fiber, seed oil, traditional and modern health benefits.

Sharma D. Phytobiomes and Their Role in Plant Disease. Shim WB, Woloshuk CP. 2001. Regulation of fumonisin B1 biosynthesis and conidiation in *Fusarium verticillioides* by a cyclin-like (C-type) gene, *FCC1*. Applied and Environmental Microbiology 67:1607–1612.

Singh VK, Singh HB, Upadhyay RS. 2017. Role of fusaric acid in the development of ‘Fusarium wilt’symptoms in tomato: Physiological, biochemical and proteomic perspectives. Plant physiology and biochemistry 118:320–332.

Smith TK, Sousadias MG. 1993. Fusaric acid content of swine feedstuffs. Journal of agricultural and food chemistry 41:2296–2298.

Song H-H, Lee H-S, Jeong J-H, Park H-S, Lee C. 2008. Diversity in beauvericin and enniatins H, I, and MK1688 by *Fusarium oxysporum* isolated from potato. International journal of food microbiology 122:296–301.

Stipanovic R, Puckhaber L, Liu J, Bell A. 2011. Phytotoxicity of fusaric acid and analogs to cotton. Toxicon 57:176–178.

Studt L, Janevska S, Niehaus EM, Burkhardt I, Arndt B, Sieber CM, Humpf HU, Dickschat JS, Tudzynski B. 2016. Two separate key enzymes and two pathway-specific transcription factors are involved in fusaric acid biosynthesis in *Fusarium fujikuroi*. Environmental Microbiology 18:936–956.

Sun X, Wang Y, Wang J, Liu Y, Song D, Fu J, Jones D, Wang D, Liu M, Ma L. 2025. Comparative genomic analysis of *Fusarium oxysporum* f. sp. *lycopersici* reveals telomeric duplications of a lineage-specific region carrying *SIX8* and *PSL1* and genome-wide expansion of Foxy transposable elements. International Journal of Biological Macromolecules 297:139636.

Tausif M, Jabbar A, Naeem MS, Basit A, Ahmad F, Cassidy T. 2018. Cotton in the new millennium: advances, economics, perceptions and problems. Textile Progress 50:1–66.

Ulloa M, Hutmacher R, Zhang J, Schramm T, Roberts PA, Ellis ML, Dever J, Wheeler TA, Witt T, Sanogo S. 2023. Registration of 17 upland cotton germplasm lines with improved resistance to Fusarium wilt race 4 and good fiber quality. Journal of Plant Registrations 17:152–163.

Ulloa M, Hutmacher RB, Percy R, Wright SD, Burke J. 2016. Registration of five Pima cotton germplasm lines (Pima SJ-FR05–pima SJ-FR09) with improved resistance to Fusarium wilt race 4 and good lint yield and fiber quality. Journal of Plant Registrations 10:154–158.

Ulloa M, Hutmacher RB, Schramm T, Ellis ML, Nichols R, Roberts PA, Wright SD. 2020a. Sources, selection and breeding of Fusarium wilt (*Fusarium oxysporum* f. sp. *vasinfectum*) race 4 (FOV4) resistance in Upland (*Gossypium hirsutum* L.) cotton. Euphytica 216:1–18.

Ulloa M, Hutmacher RB, Schramm T, Ellis ML, Nichols R, Roberts PA, Wright SD. 2020b. Sources, selection and breeding of Fusarium wilt (*Fusarium oxysporum* f. sp. *vasinfectum*) race 4 (FOV4) resistance in Upland (*Gossypium hirsutum* L.) cotton. Euphytica 216:109.

USDA-ERS. 2022. Cotton sector at a glance. Department of Agriculture, Economic Research Service.

Wagner TA, Bell AA, Castles ZA, Ali A, Flores O, Liu J. 2022. Detection, genotyping and virulence characterization of Fusarium wilt race 4 (VCG0114) causing cotton wilt in three Texas fields. Journal of Phytopathology 170:492–503.

Wang Q, Cobine PA, Coleman JJ. 2018. Efficient genome editing in *Fusarium oxysporum* based on CRISPR/Cas9 ribonucleoprotein complexes. Fungal Genetics and Biology 117:21–29.

Yang P, Chen Y, Wu H, Fang W, Liang Q, Zheng Y, Olsson S, Zhang D, Zhou J, Wang Z. 2018. The 5-oxoprolinase is required for conidiation, sexual reproduction, virulence and deoxynivalenol production of Fusarium graminearum. Current genetics 64:285–301.

Zhang H. 2017. Characterization of Fsr1-Interacting Complex and its downstream pathogenic subnetwork modules in Fusarium verticillioides.

Zhang H, Kim MS, Huang J, Yan H, Yang T, Song L, Yu W, Shim WB. 2022. Transcriptome analysis of maize pathogen *Fusarium verticillioides* revealed FvLcp1, a secreted protein with type-D fungal LysM and chitin-binding domains, that plays important roles in pathogenesis and mycotoxin production. Microbiological Research 265:127195.

Zhang H, Mukherjee M, Kim JE, Yu W, Shim WB. 2018. Fsr1, a striatin homologue, forms an endomembrane-associated complex that regulates virulence in the maize pathogen *Fusarium verticillioides*. Molecular plant pathology 19:812–826.

Zhang H, Yan H, Shim WB. 2019. *Fusarium verticillioides* SNARE protein FvSyn1 harbours two key functional motifs that play selective roles in fungal development and virulence. Microbiology 165:1075–1085.

Zhang J, Lu J, Zhu Y, Huang Q, Qin L, Zhu B. 2022. Rhizosphere microorganisms of *Crocus sativus* as antagonists against pathogenic Fusarium oxysporum. Frontiers in Plant Science 13:1045147.

Zhang Y, Ma L-J. 2017. Deciphering pathogenicity of *Fusarium oxysporum* from a phylogenomics perspective. Advances in genetics 100:179–209.

Zhu Y. 2022. Fusarium wilt in cotton caused by Fusarium oxysporum f. sp. vasinfectum race 4: characterization, pathogenicity, infection process, resistance evaluation techniques, and quantitative trait locus mapping of resistance. New Mexico State University.

Zhu Y, Abdelraheem A, Cooke P, Wheeler T, Dever JK, Wedegaertner T, Hake K, Zhang J. 2022. Comparative analysis of infection process in Pima cotton differing in resistance to Fusarium wilt caused by *Fusarium oxysporum* f. sp. *vasinfectum* race 4. Phytopathology 112:852–861.

Zhu Y, Elkins-Arce H, Wheeler TA, Dever J, Whitelock D, Hake K, Wedegaertner T, Zhang J. 2023. Effect of growth stage, cultivar, and root wounding on disease development in cotton caused by Fusarium wilt race 4 (*Fusarium oxysporum* f. sp. *vasinfectum*). Crop Science 63:101–114.

Zhu Y, Lujan P, Wedegaertner T, Nichols R, Abdelraheem A, Zhang J, Sanogo S. 2020. First report of *Fusarium oxysporum* f. sp. *vasinfectum* race 4 causing Fusarium wilt of cotton in New Mexico, USA. Plant Disease 104:588–588.

